# Recurrent alternative splicing isoform switches in tumor samples provide novel signatures of cancer

**DOI:** 10.1101/006908

**Authors:** Endre Sebestyén, Michał Zawisza, Eduardo Eyras

**Affiliations:** Computational Genomics, Universitat Pompeu Fabra, Dr. Aiguader 88, E08003 Barcelona, Spain; Universitat Politècnica de Catalunya, Jordi Girona 1-3, Barcelona E08034, Spain; Catalan Institution for Research and Advanced Studies, Passeig Lluís Companys 23, E08010 Barcelona, Spain

## Abstract

Cancer genomics has been instrumental to determine the genetic alterations that are predictive of various tumor conditions. However, the majority of these alterations occur at low frequencies, motivating the need to expand the catalogue of cancer signatures. Alternative pre-mRNA splicing alterations, which bear major importance for the understanding of cancer, have not been exhaustively studied yet in the context of recent cancer genome projects. In this article we analyze RNA sequencing data for more than 4000 samples from The Cancer Genome Atlas (TCGA) project, including paired normal samples, to detect recurrent alternative splicing isoform switches in 9 different cancer types. We first investigate whether alternative splicing isoform changes are predictive of tumors by applying a rank-based algorithm based on the reversal of the relative expression of transcript isoforms. We find that consistent alternative splicing isoform changes can separate with high accuracy tumor and normal samples, as well as some cancer subtypes. We then searched for those changes that occur in the most abundant isoform, i.e isoform switches, and are therefore more likely to have a functional impact. In total we detected 244 isoform switches, which are associated to functional pathways that are frequently altered in cancer and also separate tumor and normal samples accurately. We further assessed whether these isoform changes are associated to somatic mutations. Surprisingly, only a few cases appear to have association, including the putative tumor suppressor FBLN2 and the tumor driver MYH11, which show association of an isoform switch to mutations and indels on the alternatively spliced exon. However, the number of observed mutations is in general not sufficient to explain the frequency of the found isoform switches, suggesting that recurrent isoform switching in cancer is mostly independent of somatic mutations. In summary, we present an effective approach to detect novel alternative splicing signatures that are predictive of tumors. Moreover, the same methodology has led to uncover recurrent isoform switches in tumors, which may provide novel prognostic and therapeutic targets.

Software and data are available at: https://bitbucket.org/regulatorygenomicsupf/iso-ktsp and http://dx.doi.org/10.6084/m9.figshare.1061917

## Introduction

Cancer genome projects are instrumental to describe the genetic heterogeneity of tumors and to uncover recurrent alterations that may serve as new biomarkers for prognosis and therapeutic targets (TCGA 2012a-d, TCGA 2013). However, known actionable alterations tend to occur at low frequency or are often absent in an individual tumor sample, which hinders the choice of appropriate therapeutic strategies (Vogelstein et al. 2013, Hudson 2013). There is therefore a need to expand the catalogue of molecular signatures in cancer. Alternative splicing alterations, which bear major importance in terms of the understanding and treatment of cancer (Bonomi et al. 2013), have not been exhaustively studied yet in the context of recent cancer genomics efforts. Alternative splicing may confer a selective advantage to the tumor, such as angiogenesis (Amin et al. 2011), proliferation (Bechara et al. 2014), cell invasion (Venables et al. 2013) and avoidance of apoptosis (Izquierdo et al. 2005). Some of these alterations may be caused by somatic mutations (Ward and Cooper 2010), but can also take place as a result of changes in expression, amplifications and deletions in splicing factors (Karni et al. 2007, Furney et al. 2013). This suggests that similar splicing alterations may have different genetic origins and still confer equivalent tumorigenic properties to cells. Accordingly, to uncover potential markers of prognosis and targets of therapy, it is of utmost relevance to describe the patterns of alternative splicing in tumors.

Numerous genome wide surveys have highlighted the role of alternative splicing patterns in tumors. These have mostly been based on the measurement of local patterns of splicing, encoded as events, and studied using microarrays (Thorsen et al. 2008, Lapuk et al. 2010, Misquitta-Ali et al. 2011), RT-PCR platforms (Klinck et al. 2008), or RNA sequencing (Liu et al. 2012). The description of alternative splicing in terms of simple events facilitates the validation using PCR methods and the characterization of regulatory mechanisms using sequence analysis and biochemical approaches. However, alternative splicing takes place through a change in the relative abundance of the transcript isoforms expressed by a gene, and splicing alterations important for tumor progression may involve complex patterns that are not easily described in terms of simple events. Accordingly, to ultimately determine the functional impact of splicing alterations, it is important to describe these in terms of transcript isoforms changes. This has been shown to be relevant for TP53, which produces multiple isoforms with complex variations and with different roles in tumors (Bourdon et al. 2005). Similarly, recent analyses have shown that there are transcript isoforms specific of lung and breast cancers (Kalari et al. 2012, Eswaran et al. 2013), and that transcript analysis can improve expression-based tumor classification (Zhang et al. 2013, Pal et al. 2014). Additionally, transcript isoform changes can be essential to detect resistance to anti-tumor drugs (Mitra et al. 2009, Poulikakos et al. 2011). Thus, the detection of transcript isoform changes characterizing specific tumor types can provide new cancer signatures and could be crucial for the development of diagnostic, prognostic and therapeutic strategies.

With the aim to describe the transcript isoform changes that are characteristic of tumors, we have analyzed more than 4000 RNA sequencing (RNA-Seq) samples available from the Cancer Genome Atlas (TCGA) project. In order to perform an analysis that is robust to the variability between samples from different individuals, we have applied a new rank-based algorithm that searches for consistent reversals of relative isoform expression. This algorithm provides the minimal set of isoform-pairs with relative expression changes that can accurately separate tumor from normal samples. Moreover, the obtained isoform-pairs can accurately classify unseen tumor data. We have applied the same algorithm to breast, lung and colon cancer subtypes to obtain isoform signatures that separate subtypes from each other. In particular, we found a highly significant signature for basal-like breast tumors that distinguish them from other breast cancer subtypes. We also found that a number of the identified significant isoform changes correspond to transcript isoform switches, for which the relative expression change occurs in the most abundant isoform, and are therefore more likely to have a functional impact. We found a total of 244 isoform switches in all cancer types. These switches can also accurately separate tumor and normal samples, and occur in genes belonging functional pathways frequently altered in cancer. Interestingly, only a few of these switches can be explained by somatic mutations in the gene locus, suggesting that recurrent isoform switching in cancer is mostly independent of somatic mutations. On the other hand, we found that for at least one case the isoform switch is mutually exclusive with mutations affecting the protein-coding region of the transcripts, suggesting that splicing alterations represent an alternative route towards cellular transformation. Our analyses show that recurrent transcript isoform switches represent important novel signatures in cancer that can serve as molecular markers and could lead to the development of new therapeutic targets.

## Results

### Systematic analysis of splicing isoform changes in cancer

Changes in the relative abundance of the alternative transcripts from a gene translate into a variation in their relative order in the ranking of transcript expression. Accordingly, the problem of finding alternative splicing changes in cancer can be reformulated in terms of the consistency of the reversals of their relative expression of transcript isoforms from the same gene. For this purpose we developed Iso-kTSP, which extends the principle of consistent expression reversals for gene expression (Geman et al. 2004, Tan et al. 2005, Price et al. 2007) to alternative splicing isoforms. The Iso-kTSP algorithm stores the ranking of isoform expression from multiple samples separated into two classes (Figure 1A). All possible isoform pairs from the same gene are sorted according to the sum of frequencies of the two possible relative orders occurring separately in each class, defined as score *S_1_* (Methods). This score provides an estimate of the probability for the isoforms to change relative order between the two classes. The top scoring isoform pairs are therefore the most consistent changes in isoform relative abundance for a gene between two classes, tumor and normal, or between two tumor subtypes. Each one of these isoform pairs provides a classification rule based on the relative expression order, with a discrimination power related to the consistency of this reversal across samples.

**Figure 1.**
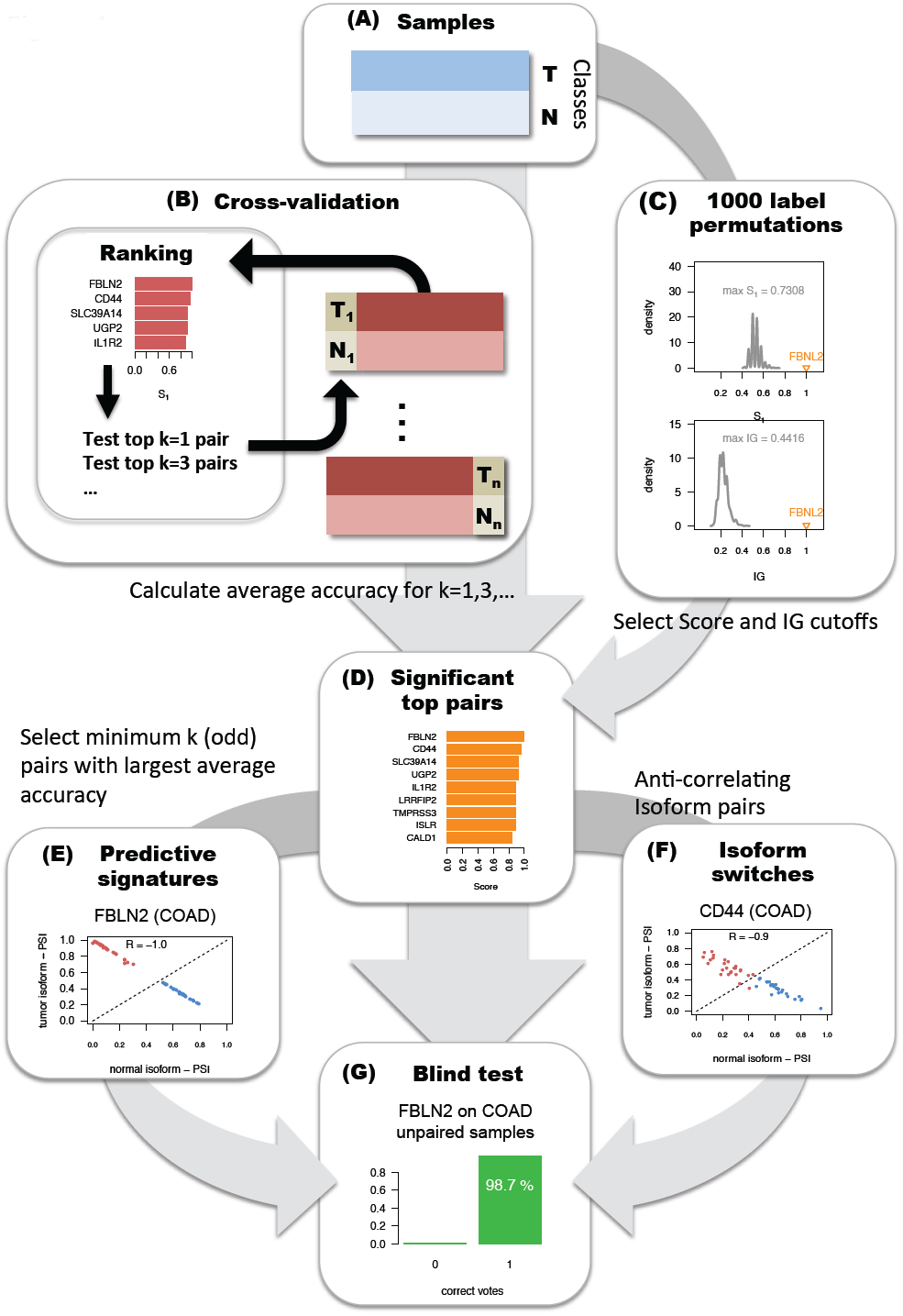
Methodology for detecting significant transcript isoform changes in cancer. The method is illustrated with data from colon adenocarcinoma (COAD). **(A)** Samples are partitioned into two groups, here tumor (T) and normal (N). **(B)** Cross-validation is performed by iteratively training in all but one pair of tumor-normal samples, and testing on this left-out sample pair. At each step of the cross-validation, the top k=1,3,5, etc. isoform-pairs according to score *S_1_* are tested on the left-out sample pair. **(C)** Significance is assessed by comparing to the expected values obtained from 1000 permutations of the class labels and by selecting at each permutation the isoform-pair with the highest score *S_1_*. **(D)** The result is a global ranking of isoform-pairs that change relative expression between tumor and normal samples more than expected by change. **(E)** A minimal classification model is obtained by selecting the smallest number of pairs from the global ranking with the largest average accuracy calculated in the cross-validation. The example shows that for colon this is a single pair model in gene FBLN2. **(F)** From the global ranking of significant isoform-pairs, we predict as isoform switches those that anti-correlate across samples. In the example, CD44 presents a clear switch between two isoforms even though it was not chosen in the minimal classification model. **(G)** The isoform-pairs (either from the minimal classification model or from the set of isoform switches) are tested on a held-out dataset of unpaired tumor samples.

Using cross-validation, the ranking of isoform-pairs is calculated at each iteration step on a balanced set leaving out one sample from each class, which are used for testing (Figure 1B). The prediction class for a new sample is obtained by evaluating the expression ranking in the new sample against the isoform pair rules. At each iteration step in the cross-validation, the top *k*-pairs (*k*=1… *k*_max_, with *k* odd) are evaluated on the test set. Each isoform-pair from the same gene defines a rule, where each gene is only used once. The rule is defined such that for a pair, if the first isoform has lower expression than the second the sample is predicted to be normal, otherwise is predicted to be a tumor. Accordingly, the first isoform is defined as the tumor isoform, and the second as the normal isoform. Significance of the isoform-pair rules is measured by performing 1000 permutations of the sample labels (Figure 1C). At each permutation, the algorithm is run as before keeping only the pair with the highest score *S_1_*. An isoform-pair is defined as significant if its score *S_1_* and information gain (IG) are larger than the maximum ones obtained from the permutation analysis. The global ranking of isoform-pairs and permutation analysis yields the list of significant isoform changes (Figure 1D). From this set, we derive a minimal classification model with the smallest odd number of isoform-pairs with the highest average performance (Figure 1E). The final classification is based on simple majority voting with an odd number of isoform pairs. On the other hand, some of the isoform-pairs from the list of significant cases are in fact isoform switches, for which the relative expression change occurs in the most abundant isoform of the gene, and which we detect by the anti-correlation of the relative inclusion levels or PSIs of the isoforms (Figure F). Finally, to further assess the accuracy of the minimal classification model, or that of a set of isoform switches, a blind test is carried out on samples that were not used for cross-validation (Figure 1G). On this set we measure the proportion of samples correctly labeled by the classifier, as well as the number of correct votes for each prediction.

### Recurrent alternative splicing isoform changes can separate tumor and normal samples

We downloaded the available RNA-Seq datasets for tumor and paired normal samples from TCGA for 9 cancer types (Table 1). Prediction of tissue types from gene expression with URSA (Lee et al. 2013) was used to detect and filter out outliers in the samples (Supplementary Figure 1). The tumor and paired normal samples that were kept show a remarkably similar pattern of tissue expression in BRCA, COAD, KIRC, PRAD and THCA (Supplementary Figure 1), and partly in KICH. On the other hand, LUAD, LUSC, and HNSC show heterogeneous patterns of predicted tissue types, probably due the mixed cellular origins (Lundin et al. 2013). The list of samples used for further analyses are given in Supplementary File 1.

**Table 1.**
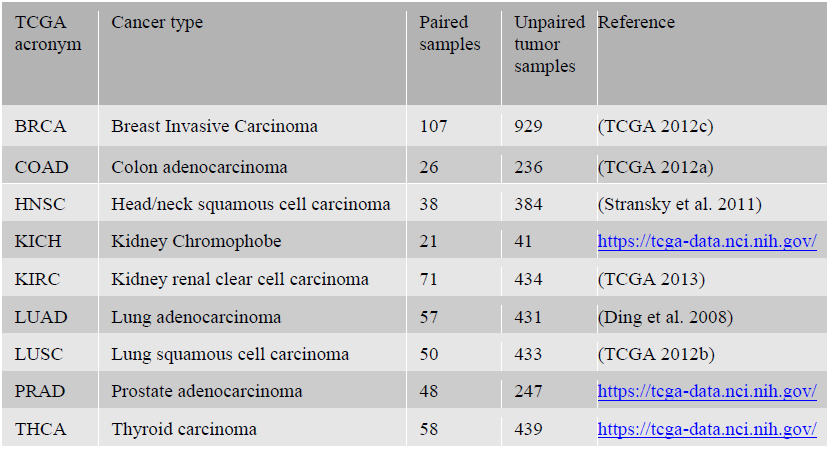
Number of analyzed paired and unpaired tumor samples from each cancer type: Breast carcinoma (BRCA), colon adenocarcinoma (COAD), head and neck squamous cell carcinoma (HNSC), kidney chromophobe (KICH), kidney renal clear-cell carcinoma (KIRC) lung adenocarcinoma (LUAD), lung squamous cell carcinoma (LUSC), prostate adenocarcinoma (PRAD), and thyroid carcinoma (THCA). All datasets were obtained from https://tcga-data.nci.nih.gov/. For the paired samples we also used the corresponding normal samples from the same patients. For the list of samples used see Supplementary File 1.

| TCGA acronym | Cancer type | Paired samples | Unpaired tumor samples | Reference |
| --- | --- | --- | --- | --- |
| BRCA | Breast Invasive Carcinoma | 107 | 929 | (TCGA 2012c) |
| COAD | Colon adenocarcinoma | 26 | 236 | (TCGA 2012a) |
| HNSC | Head/neck squamous cell carcinoma | 38 | 384 | (Stransky et al. 2011) |
| KICH | Kidney Chromophobe | 21 | 41 | https://tcga-data.nci.nih.gov/ |
| KIRC | Kidney renal clear cell carcinoma | 71 | 434 | (TCGA 2013) |
| LUAD | Lung adenocarcinoma | 57 | 431 | (Ding et al. 2008) |
| LUSC | Lung squamous cell carcinoma | 50 | 433 | (TCGA 2012b) |
| PRAD | Prostate adenocarcinoma | 48 | 247 | https://tcga-data.nci.nih.gov/ |
| THCA | Thyroid carcinoma | 58 | 439 | https://tcga-data.nci.nih.gov/ |

For each cancer type, the Iso-kTSP algorithm was applied to the paired samples to obtain minimal classifiers to separate tumor and normal samples, and the blind test performed on the remaining unpaired tumor samples. This yielded different predictive models for the 9 cancer types (Figure 2A and Supplementary Figure 4A), with PRAD, THCA and KIRC having the lowest average accuracies, and LUSC, LUAD, COAD and KICH having 100% average accuracy. Moreover, the blind tests of these models on the unpaired tumor samples show overall accuracies greater than 84% (Figure 2B and Supplementary Figure 4B). These models provide a minimal set of isoform pairs whose relative expression can separate tumor and normal samples with high accuracy despite the variability of the transcript expression measurement across samples (Supplementary Figures 3-8) (Model files are given in Supplementary File 2). All the isoform-pairs in the derived models are significant according to our permutation analysis (Supplementary Figure 9). This significance also depends in general on the number of samples available and on the heterogeneity of the tumor samples. Permutation analysis for a varying number of input samples indicates that in order to obtain significant isoform changes, more than 13 samples are needed on average (Supplementary Figure 10), which is fulfilled by the cancer types analyzed.

The isoform changes detected include FBLN2, which appears as a single gene model for COAD and is part of the BRCA model (Figure 3A). FBLN2 has been proposed before to be a tumor suppressor (Law et al. 2012) and its cancer-related function seems to be specific of the protein produced in tumor cells (Baird et al. 2013). We found that FBLN2 undergoes an isoform change related to the skipping of a protein coding exon (Supplementary Figure 11). Additionally, this isoform switch occurs in more than 98% of the unpaired tumor samples in BRCA and COAD (Figure 1A). In the case of LUAD, surprisingly, we found that the most informative isoform change does not occur in NUMB, known to have a splicing switch related to proliferation (Misquitta-Ali et al. 2011, Bechara et al. 2014), but in QKI. In fact, the isoform change in QKI cannot be described in terms of a simple alternative splicing event (Figure 2C). In contrast, LUSC model involves a different set of genes compared to LUAD, including the gene ZNF385A (Figure 2), whose protein product interacts with TP53 thereby promoting growth arrest (Das et al. 2007). The isoform change found is related to the use of an alternative first exon and alternative 3’ splice-site (Supplementary Figure 12). Similarly to COAD, THCA and KIRC have single-gene models (Supplementary Figure 2). In particular, for THCA the model involves S100A13, which codes for a calcium binding protein and which has been proposed to be a new marker of angiogenesis in various cancer types (Massi et al. 2010). The isoform change involves an alternative first exon and classifies correctly 84.5% of the unpaired tumor samples (Supplementary Figure 13). Interestingly, S100A13 and another member of the S100 family, S100A16, also have isoform changes in KICH, even though they were not included in the KICH model (Supplementary Figure 13). The single isoform change in KIRC involves the production of an intron-retention transcript, annotated as non-coding in the gene CPAMD8 (Supplementary Figure 14). A similar case occurs in the gene NAGS, which is part of the KICH model (Supplementary Figure 2) and is related to an autosomal recessive urea cycle disorder (Häberle et al. 2003). We predict that a protein coding isoform changes in tumors into an isoform with a retained-intron that is annotated as non-coding (Supplementary Figure 15). Importantly, the loss of the protein coding isoform is predictive of 100% of the KICH tumor samples (Supplementary Figure 15). Other isoform changes are discussed in the supplementary material (Supplementary Figures 16-18). GFF tracks for the isoform-pairs in all derived models can be found in Supplementary File 3.

**Figure 2.**
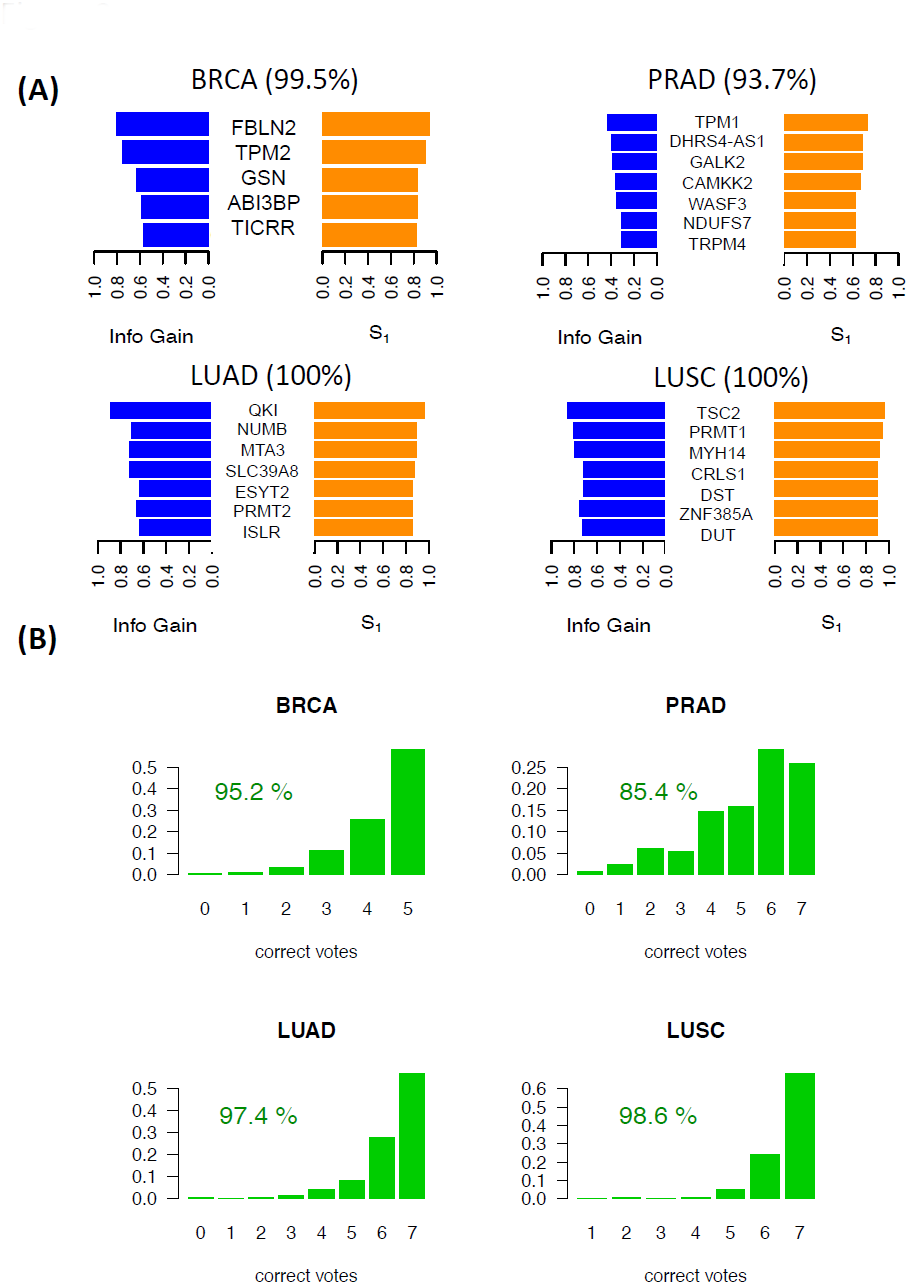
Predictive isoform-pair models. **(A)** Minimal isoform-pair classifiers for BRCA, PRAD, LUAD and LUSC (models for KICH, KIRC, HNSC and THCA are given in Supplementary Figure 2). Each panel shows the score *S_1_* and information gain (IG) for each isoform-pair in the model. All the isoform-pairs are significant according to the permutation analysis. Next to each cancer label the maximum expected accuracy is given, which is calculated from the cross-validation analysis. Plots of the expression values for each isoform pair are provided in Supplementary Figures 3-8. **(B)** Blind tests of the isoform-pair models on the unpaired samples for each cancer type. The barplots indicate the proportion of samples (y-axis) for each number of possible correct votes (x-axis), i.e., the number of isoform-pair rules from the model fulfilled by the tumor sample. A sample is labeled according to a majority vote from all isoform-pair rules.

**Figure 3.**
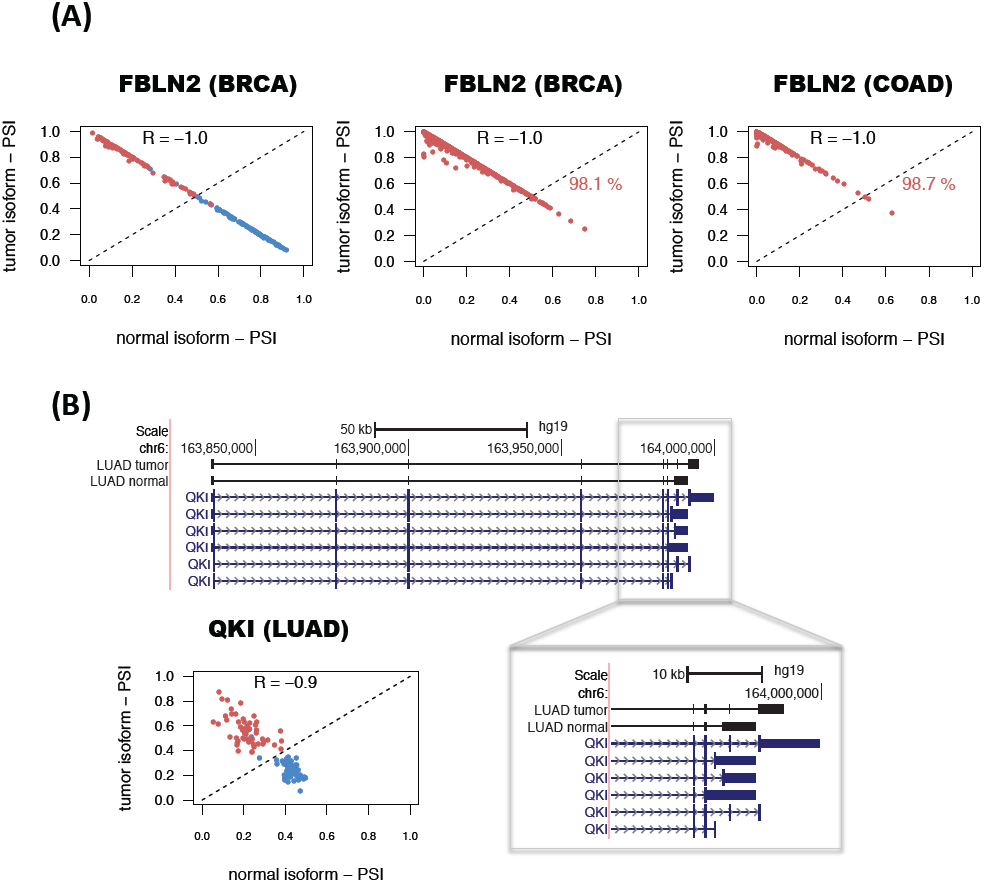
Examples of predictive isoform-pairs. **(A)** Relative inclusion values (PSIs) for the isoform-pair in FBLN2 predicted to separate well tumor from normal in BRCA and COAD. The x-axis represents the inclusion level values (PSI) for the normal isoform and the y-axis the value for the tumor isoform. Tumor samples are shown in red, whereas normal samples are shown in blue. The left panel shows the PSIs in paired samples, whereas the middle and right panels show these for the unpaired samples **(B)** Isoform change for QKI in LUAD samples. The gene locus of QKI is shown, indicating the exon-intron structures of the isoforms predicted to be the tumor and normal isoforms for LUAD. The zoom-in highlights the 3’ end region where the splicing variation takes place. The bottom left panel shows the PSI values for the normal (x-axis) and tumor isoforms (y-axis) for the normal (blue) and tumor (red) paired samples.

## Changes in alternative splicing isoforms can discriminate tumor subtypes

Cancers are generally classified into subtypes to facilitate patient stratification for more precise prognosis and selection of therapeutic strategy. In particular, breast cancer classification has been recently refined based on molecular information from multiple sources (TCGA 2012c). We thus decided to investigate whether breast cancer subtypes are associated with consistent isoform changes when compared to each other. We separated the BRCA tumor samples into luminal A, luminal B, Her2+ and basal-like as labeled by TCGA (TCGA 2012c) (Supplementary File 1) and run the Iso-kTSP algorithm comparing each subtype against a pool from the rest. In order to maintain balanced sets for the comparison and avoid biases due to sample selection, we subsampled 100 times 45 arbitrary samples for a given subtype and a pool of 15 from each of the other three subtypes together. At each iteration step, we performed permutation analysis of the labels to determine the significance of the detected isoform changes. We found that only basal-like tumors showed isoform changes that were significant in more than 80% of the sampling iterations (Figure 4A and Supplementary Figure 19). Among the most significant cases we found KIF1B, which has been implicated in apoptosis (Schlisio et al. 2008); ATP1A1, proposed to have tumor suppressor activity (Cao et al. 1997); ITGA6, found to be required for the growth and survival of a stem cell like subpopulation of MCF7 cells (Cariati et al. 2008); and CTNND1, whose alternative splicing was previously related to cell invasion and metastasis (Yanagisawa et al. 2008) (Figure 4A). We selected the top 7 isoform-pairs in basal-like that were significant in more than 80% of the iterations. This model classified correctly 93.6% of all the BRCA tumor samples, with 47% of the samples fulfilling all 7 isoform change rules (Figure 4B). Although this cannot be considered a blind test, it provides an estimation of the expected accuracy. For the other BRCA subtypes we found much lower consistency of the isoform changes and none of them were significant for more than 13% of the permutation tests (Supplementary Figure 20).

**Figure 4.**
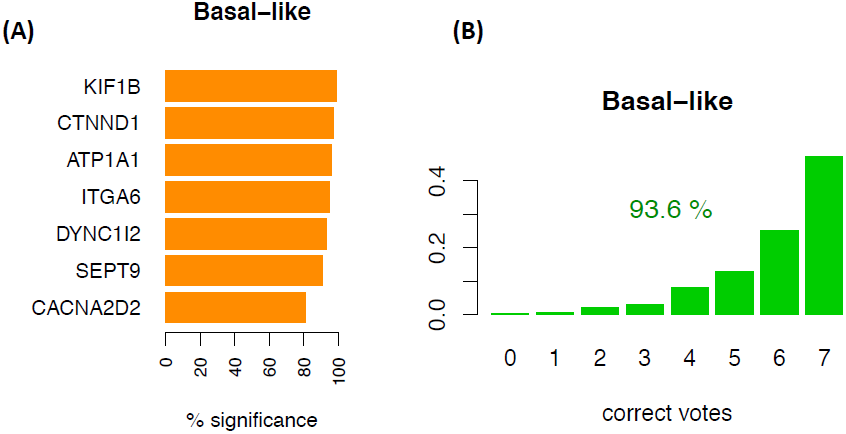
Isoform-pair rules for the basal-like breast tumors. **(A)** The top 7 recurrent isoform changes found in the 100 comparisons of subsets of basal-like samples against a balanced pool set of the other subtypes (luminal A, luminal B and Her2+). The barplots indicate the frequency of iterations for which the isoform pair was significant according to the permutation analysis performed on the same subsampled sets. All these isoform-pairs are significant according to our permutation analysis. **(B)** Accuracy of the model for the classification of basal-like samples against other subtypes when tested on the entire set of 1036 BRCA tumor samples. The barplots show the proportion of samples (y-axis) with each possible number of correct votes (x-axis), from 0 to the number of genes in the model, and the percentage of all the samples correctly classified.

Four different subtypes have been defined based on mRNA expression for the lung squamous cell carcinomas (LUSC): basal, classical, primitive and secretory; which have different clinical and biological characteristics (Wilkerson et al. 2010). We applied the same approach as above to the four LUSC subtypes using the subtype labeling from TCGA (TCGA 2012b), comparing 24 samples from each subtype against the pool of three sets of 8 arbitrary samples from the other subtypes. The most relevant isoform change was found for gene GCNT2 in association to the classical subtype in at least 60% of the subsampling iterations, but significant only in 22% of them (Supplementary Figure 21 and 22). Interestingly, GCNT2 overexpression has been linked to breast and lung cancer metastasis (Zhang et al. 2011). All other found changes occurred at lower frequencies and showed significance in no more than 3% of the iterations. Despite of the low recurrence of the isoform changes, tests on the entire set of LUSC tumor samples was able to separate correctly classical from other subtypes for more than 80% of them (Supplementary Figure 21).

Colorectal cancers have been classified into hypermutated and non-hypermutated, where non-hypermutated tumors have generally worse prognosis (TCGA 2012a). Following previous definitions (TCGA 2012a), we labeled COAD samples as hypermutated if they had more than a total of 250 mutations, and as non-hypermutated those with less than 250 mutations. We then compared both subtypes by subsampling 40 samples from each one 100 times. This analysis yielded specific isoform changes between the two types occurring in more than 40% of the iterations (Supplementary Figure 23), including a change in the long non-coding RNA gene antisense of NUTM2A (NUTMA2A-AS1), which appeared in 57% of the models. We tested two different models with the top 5 and 13 isoform-pairs, obtaining an accuracy of more than 80% on the total COAD dataset (Supplementary Figure 18). Models and GFF tracks for all subtype models are provided in Supplementary Files 2 and 3, respectively.

### A catalogue of alternative isoform switches in cancer

The alternative isoform changes described above can separate tumor and normal samples, and in some cases specific cancer subtypes. However, these models are optimized to have the minimum number of isoform-pairs and maximum average accuracy, which is convenient for defining biomarkers with potential clinical applications. On the other hand, the frequency of these isoform changes does not imply functional relevance. However, among the recurrent isoform-changes, those more likely to be functionally relevant will be the ones for which the change occurs in the most abundant isoform, i.e. isoform switches (Figure 1F). Accordingly, in order to obtain all the significant isoform changes with a possible functional relevance in cancer, we decided to calculate all the significant isoform switches between tumor and normal samples. We first retrieved all those genes with significant isoform changes according to our permutation analysis (Figure 1D). This yielded a total 1178 genes for the 9 cancer types. We further filtered these genes by imposing a score *S_1_* > 0.5, which corresponds to selecting isoform-pairs with a change in more than 75% of the samples. To select for switches, we kept those cases for which the relative inclusion levels of the isoforms anti-correlate (Figure 1F), as observed for FBLN2, QKI and other genes (Figure 3 and Supplementary Figures 12-18). We thus selected those isoform-pairs having an anti-correlation of PSI values of R < −0.8 (Spearman). Finally, we kept only those with average expression per isoform of > 1 TPMs in either tumor or normal samples.

These criteria gave rise to a total of 244 isoform switches, with 59 of them appearing in more than one cancer type (Figure 5) (Supplementary File 4). The most common of the switches is the one described above for FBLN2. From the total 244 switches, 10 occur in known cancer drivers (Figure 5). Moreover, we also found switches in genes whose splicing has been associated before with cancer, like CD44, which has been observed to be relevant in colon cancer initiation (Du et al. 2008), and SLC39A14, whose alternative splicing is regulated by WNT in colon cancer (Thorsen et al. 2011). LUAD, KIRC and LUSC are the cancer types with most switches, with 85, 65 and 54, respectively. LUSC and LUAD have 33 switches in common. In contrast, KIRC and KICH have only 2 switches in common. HNSC and PRAD are the cancer types with the fewest switches, 7 and 2 respectively. Although functional analysis did not yield any significantly enriched Reactome pathways (Croft et al. 2014), isoform switches appear frequently in signal transduction, immune system and metabolism related pathways (Figure 4 and Supplementary Figure 24). On the other hand, Gene-Ontology analysis shows enrichment of multiple categories, including actin activity in relation to cell motility and migration, in categories related to extracellular organization, as well as in response to estrogen and regulation of MAPK activity (Supplementary Figure 25).

**Figure 5.**
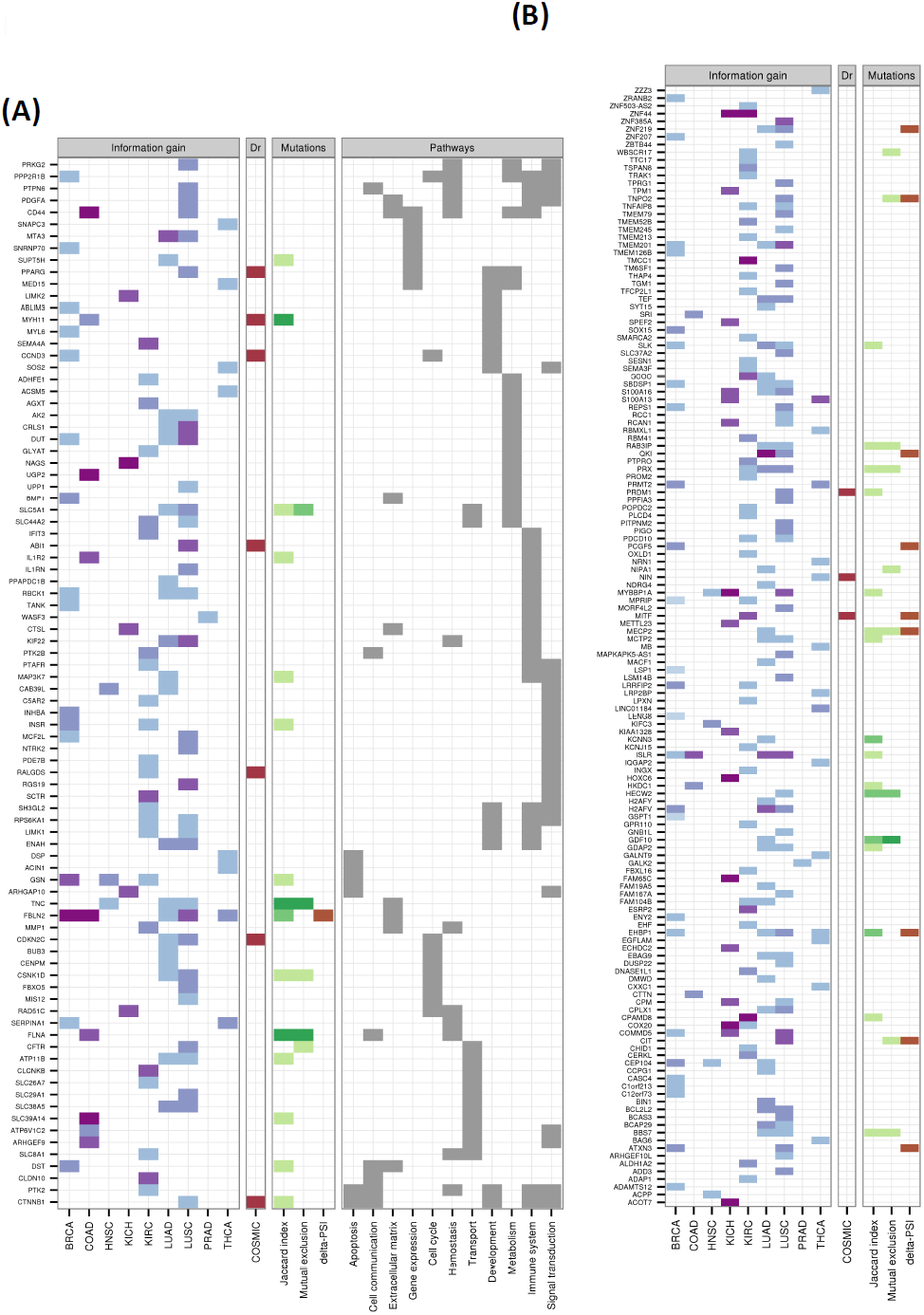
Catalogue of isoform switches across various cancer types. Heatmap of the 244 isoform switches detected for the 9 cancer types, separated according to whether the genes had an annotated Reactome pathway **(A)** or not **(B)**. The heatmaps show whether the isoform switch occurs in each cancer type, with the color code indicating the information gain (IG) of the switch: from light blue for low IG (0-0.2) to dark purple for high IG (0.8-1). In red we indicate whether the gene with the switch is annotated in COSMIC (Forbes et al. 2011) as a tumor driver. Regarding the mutations, we indicate the Jaccard index and the mutual-exclusion score with light green (0.01-0.02), medium green (0.02-0.03) and dark green (larger than 0.03). We also indicate the presence of a significant difference (p-value < 0.05) of the relative inclusion (PSI) difference between tumor and normal isoforms in mutated and non-mutated tumor samples before multiple-testing correction (brown color). We further show the Reactome Pathway annotation for those genes for which this was available.

We also tested the accuracy of switches as predictive models by performing blind tests on the unpaired tumor samples (Figure 1G) and found accuracies of around 90% and higher (Supplementary Figure 26). These isoform switches are thus good predictors of tumor samples. All details for the found isoform switches and corresponding GFF tracks are provide in Supplementary Files 4 and 5.

### Isoform switches in cancer are not frequently associated with somatic mutations

As splicing changes may be triggered by somatic mutations (Ward and Cooper 2010), we thus investigated whether any of the detected isoform switches may be caused by recurrent somatic mutations in the same genomic locus. To this end, we tested whether in tumor samples there was any association between the presence of the isoform switch and somatic mutations in the region of the transcript isoforms undergoing the switch. Since in addition to intronic mutations, synonymous as well as non-synonymous mutations could alter the splicing of a gene (Sterne-Weiler & Sanford 2014), we considered all mutation types available in TCGA: coding-related (nonsense, missense, frameshift and indel) and non-coding-related (synonymous, splice-site and RNA) mutations. For each isoform-switch and for each cancer type, we calculated the Jaccard index across all samples for the association between the presence of the switch and the presence of somatic mutations (Figure 6A) (see Methods for details). These Jaccard indexes agree with mutual information and do not correlate with the average mRNA length of the switches (Supplementary Figure 27). This analysis shows genes FBLN2, MYH11, FLNA and TNC to have the strongest association between mutations and switches (Figure 6A and Supplementary Figure 20). These four genes are also the ones with the highest frequency of mutated samples (Figure 6A). For FBLN2, we found several mutations in BRCA and COAD samples on the alternative exon and the flanking constitutive exons (Figure 6B). However, there are not enough mutations to explain all the switches observed. We also found frequent mutations in the alternatively spliced region of the oncogene MYH11. In particular, we found recurrent deletions and insertions on the alternative exon in COAD and BRCA tumor samples that coincide with the presence of the switch (Figure 6C). The location of these indels coincides with a region of low conservation and is next to a putative binding site for the splicing factor SRSF1 (Supplementary Figure 28). Nonetheless, the number of found mutations cannot explain either the frequency of the switches observed.

**Figure 6.**
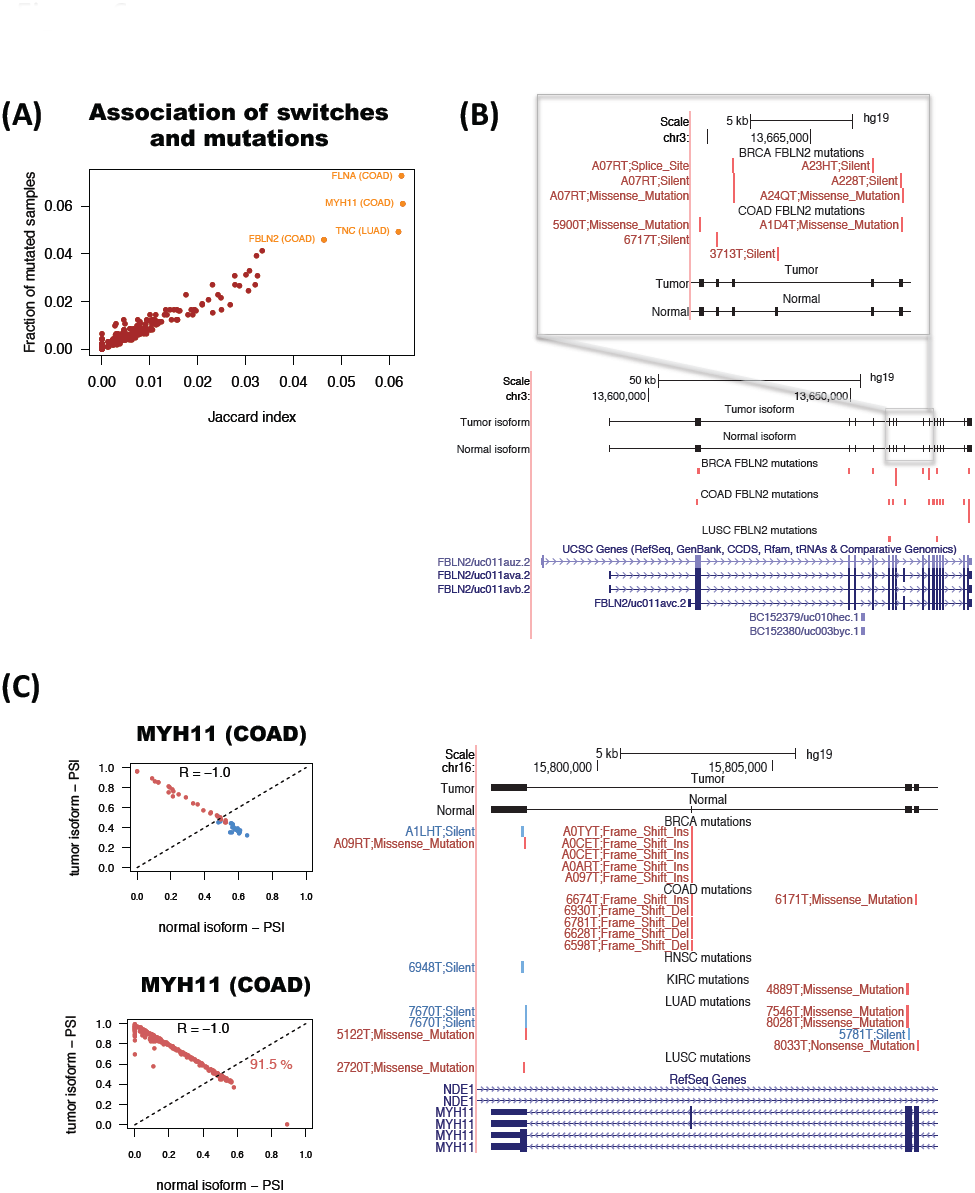
Association between somatic mutations and isoform switches. **(A)** Jaccard index (x-axis) for the association of mutations with switches in tumor samples, and frequency of samples with mutations in the transcripts undergoing the switch (y-axis). **(B)** Example of the tumor suppressor FBLN2. Mutations present in each cancer type are represented in red if the switch is present in the same sample, and blue if that sample does not have the switch. Each mutation is labeled with an identifier of the sample and the type of mutation. **(C)** Example of the oncogene MYH11. The relative inclusion values (PSI) of the two isoforms in the switch (left panels) separate tumor and normal in COAD and can classify correctly 91.5% of the unpaired tumor samples. Mutations present in each cancer type (right panel) are represented in red if the switch is present in the same sample, and blue if that sample does not have the switch. Each mutation is labeled with an identifier of the sample and the type of mutation.

Somatic mutations could also affect the magnitude of the splicing change in specific samples. We therefore tested, for each isoform-switch, whether the presence of mutations is associated with a larger difference of PSI between the tumor and normal isoforms involved in the switch (Methods). Among the four most significant cases, we found FBLN2 and EHBP1 (Supplementary Figure 29). These two cases show some differences in the distributions of samples with and without mutations (Supplementary Figure 29). However, the proportion of mutated samples is very small to make a reliable comparison and after multiple-testing correction, none of the found cases remained significant. This suggests that, except for a limited number of cases, mutations may not be the main cause of the recurrent splicing switches we have found in cancer.

We thus hypothesized that mutations and isoform switches may occur independently as two alternative mechanisms of functional transformation in cancer. To test this possibility, we measured how frequently mutations that affect the protein-coding region occur in tumor samples without the isoform switch in the same gene by defining a mutual-exclusion score based on the number of samples with no switch but with protein-affecting mutations (Methods). We found that in general the mutual exclusion score correlates with the overall proportion of mutated samples (Supplementary Figure 30). However, the number of samples with switch and mutation is generally comparable or higher, except for the gene Tenascin C (TNC), for which we find more samples with a protein-affecting mutation and no switch than with switch and protein-affecting mutation (Supplementary Figure 30). This indicates that there is no strong bias towards this mutual exclusion. We conclude that there are currently not a sufficient number of mutations that can provide an explanation of the described recurrent isoform switches. Nonetheless, there are a few cases for which this association may exist, as described for the genes FBLN2, MYH11 and TNC.

## Discussion

We have applied the principle of relative expression reversals (Geman et al. 2009, Tan et al. 2005) to the search of recurrent alternative splicing isoform changes in tumors using available RNA-Seq data from the TCGA project for 9 different cancer types. In our implementation of this algorithm for isoforms (available at: https://bitbucket.org/regulatorygenomicsupf/iso-ktsp) each classification rule is described in terms of the relative expression of a single pair of isoforms per gene. In this context, the principle of reversals has a natural interpretation as an alternative splicing change between two conditions. This algorithm provides robust classifiers despite of the variability of isoform expression across tumor samples, as the models are not dependent on parameterizations or on any normalization that would maintain the order of the isoform expression. This rank-base method is especially useful for isoform expression from RNA-Seq data, since between-sample normalization methods are not yet fully established. In fact, our approach is applicable to integrate data from heterogeneous platforms, as long as they provide a meaningful ranking of expression. Moreover, the method produces significant isoform changes from a relatively small number of samples, which makes it useful for medium-sized sequencing projects.

We have derived classification rules based on isoform changes that can distinguish tumor from normal samples, and between some tumor subtypes. The predictive models show overall a high accuracy when tested on held-out datasets. Except for the cases of colon, thyroid and kidney carcinomas, individual isoform-pair rules do not show in general a strong predictive power. However, in combination, they accurately classify tumor samples in the blind test. This suggests that splicing alterations are heterogeneous in tumor samples, but in combination they provide characteristic signatures, similarly to the patterns of somatic mutations (Vogelstein et al. 2013). This heterogeneity is further highlighted by the fact that different cancer types share a small fraction of the found isoform changes. Although some of these changes may be explained by the differences in the cell composition of tumors (Venables et al. 2013, Mallinjoud et al. 2014), we observed an homogenous pattern of predicted tissue types in tumor and normal samples for most of the cancer types analyzed, suggesting that the splicing changes are not a consequence of differences in cell type composition.

Comparative analysis between cancer subtypes only yielded a significant model for basal-like breast tumors, which includes genes with known functional relation to cancer. For the LUSC subtypes we did not find significant changes, and only the classical subtype showed frequent isoform changes in the gene GCNT2. Similarly, COAD subtypes, based on the mutations of the samples, did not show significant isoform changes, but we observed frequent changes in a non-coding RNA gene antisense of NUTMA2A. These results indicate that most of the subtypes considered have similar alternative splicing patterns.

Our initial analysis shows that isoform changes hold sufficient information to separate tumor and normal samples, and specific tumor subtypes. This suggests they could serve as effective molecular markers, as they would only require measuring the expression of two isoforms per gene, for a small number of genes. Furthermore, the application of this method to separate tumors according to clinical information will provide a useful prognostic tool. On the other hand, these signatures are not necessarily related to a biological effect specifically relevant for the tumor. To investigate this aspect, we selected all those significant isoform changes that are also isoform switches, i.e. the change occurs in the most abundant isoform, and therefore more likely to have a functional impact. We found 244 such isoform switches, which can also accurately separate tumor and normal samples, and some of which appear in multiple cancer types. Interestingly, many of the found isoform switches occur in pathways that are often altered in tumors, and some of them occur in known cancer driver genes, including CDKN2C, CTNNB1, ABI1 and MYH11. Additionally, we found that of the splicing switches cannot be described in terms of simple events. This is the case for QKI, for which we predict an isoform switch specific to lung adenocarcinoma. The found isoform switches provide an opportunity to develop experimental strategies based on the detection of specific protein isoforms. In particular, we found genes with isoform switches involved in cell communication pathways, including DST and FLNA, which could be potentially used for diagnostic or prognostic applications, or even for developing tumor-specific therapeutic targets with reduced cross-reactivity to other proteins.

Surprisingly, we did not find strong associations of somatic mutations with the isoform switches. It has been recently proposed that synonymous mutations in known cancer drivers may contribute to the oncogenic process (Supek et al. 2014). However, a direct link was not made between the observed somatic mutations and specific splicing changes measured in the same tumor samples. We found only a handful of genes with significant association between the isoform change and somatic mutations occurring in the same samples. These include the putative tumor suppressor FBLN2 and the cancer driver MYH11, the latter showing a recurrent indel in the alternative exon in samples where it is skipped. Still, mutations alone cannot explain the splicing changes observed, as 99% of the transcripts analyzed appear mutated in less than 5% of the tumor samples, whereas the majority of switches occur in at least 50% of the samples. This could mean that there are many intronic mutations not represented in the currently available exome-based data, which could explain the observed variations. Alternatively, the recurrent switches could be explained by alterations in splicing factors. Although point mutations and indels on splicing factors do not occur with sufficient frequency to explain the switches (Furney et al. 2013), some splicing factors show frequent amplifications, deletions and changes in expression in tumors (Karni et al. 2007). Another interesting hypothesis is whether alterations in chromatin modifications or DNA methylation may be responsible for the observed changes. These alterations are frequent in cancers (Esteller 2007, Ellis et al. 2008) and can induce changes in splicing (Luco et al. 2010, Maunakea et al. 2013). Interestingly, the gene FBLN2, which presents a switch in various cancers, has been observed frequently methylated in breast and other epithelial tumors (Hill et al. 2010).

Consistent isoform switching in tumors thus seems generally independent of somatic mutations; and moreover, only 10 of the detected 244 switches occur in known cancer drivers. This raised the question of whether these two alterations could be actually mutually exclusive. We found that only TNC, linked to cell invasion in tumors (Hancox et al. 2009), has this mutual exclusion pattern between the isoform switch and somatic mutations affecting the coding regions, suggesting that, albeit to a limited extent, splicing switches may provide an alternative mechanism towards functional transformation in cancer. In summary, we have detected recurrent alternative splicing isoform changes that are predictive of various tumoral conditions, and which may have potential applications for diagnostic and prognostic purposes. The same methodology has allowed us to uncover recurrent isoform switches in tumors, which are likely to have a functional impact, and which may be useful to explore novel therapeutic strategies. Further research will be necessary to determine the functional impact produced by the described isoform changes and how these may actually contribute to the tumor. We hypothesize that the observed recurrent changes in splicing, regardless of their cause, may contribute together with mutations and other alterations to explain tumor formation; hence, providing novel signatures for cancer.

## Methods

Available processed RNA-Seq data for tumor and normal samples was downloaded from the TCGA data portal (https://tcga-data.nci.nih.gov/tcga/) for all cancer types together with the UCSC gene annotation from June 2011 (assembly hg19) and the somatic mutation data. To assess sample quality, the provided estimated read-counts per gene were analyzed using URSA (Lee et al. 2013) and sample pairs that did not cluster with the rest of the samples were removed (Supplementary Figure 1). The list of kept samples can be found in Supplementary File 1.

The abundance of every transcript per sample was calculated in transcripts per million (TPM) (Li et al. 2010) from the transcript-estimated read counts provided by TCGA and the isoform lengths from the UCSC (June 2011) annotation. No further normalization on the TPM values was performed. For each transcript, the relative abundance (or PSI) per sample was calculated by normalizing the TPM by the sum of TPMs for all transcripts in the gene. Genes with one single isoform or no HUGO ID were removed for the Iso-kTSP analysis.

The Iso-kTSP algorithm is based on the kTSP algorithm (Geman et al. 2004, Tan et al. 2005). It stores the isoform expression rankings in samples from two groups, *C_m_, m=1,2*. For every pair of isoforms *I_g,i_* and *I_g,j_* in each gene *g*, Iso-kTSP calculates a score based on the frequencies of the two possible relative orders in both classes:

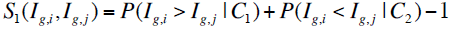

Where *P(I_g,i_* > *I_g,j_* |*C_1_)* and *P(I_g,i_* < *I_g,j_* |*C_2_)* are the frequencies at which the isoform *I_g,i_* appears later than, or before, *I_g,j_* in the expression ranking of classes *C_1_* or *C_2_*, respectively. To avoid possible ties, a second score *S_2_* is used, which is based on the average rank difference per class *C_m_* for each isoform pair, as proposed previously (Tan et al. 2005). Defining *R(I_g,i_ |S_a_,C_m_)* as the rank of isoform *I_g,i_* in sample *S_a_* and class *C_m_*, the average rank difference between two isoforms in a given class is calculated as

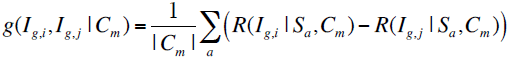

where |*C_m_*| denotes the number of samples in class *C_m_*. The score *S_2_* for an isoform-pair is then defined as (Tan et al. 2005):

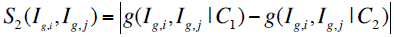

All possible isoform pairs are then sorted by the *S_1_* score and in the case of a tie, by the *S_2_* score. Moreover, only pairs of isoforms from the same gene are considered and only a single pair of isoforms per gene is listed in a ranking of isoform-pairs. Classification rules are given in terms of *k* isoform pairs. The classification of a new sample is performed by evaluating each isoform-pair rule against the ranking of isoform expression of the new sample. For each isoform-pair rule, the classifier selects the class for which the data fulfills the rule. The final decision for classification is established by simple majority voting, by selecting the most voted class from the *k* rules. In order to avoid ties in the voting, *k* is always odd. Significance of the computed isoform changes was evaluated by shuffling labels from the two classes 1000 times. For each shuffling step, the Iso-kTSP algorithm was re-run and the top-scoring isoform-pair was selected. An isoform-pair is defined as significant if its Information Gain and Score *S_1_* are larger than any of the values obtained from the 1000 shufflings of the same cancer type. The Iso-kTSP is implemented in Java. Software and documentation are available at https://bitbucket.org/regulatorygenomicsupf/iso-ktsp.

For the purpose of finding associations, the number of samples for which a gene has a switch was compared with the number of samples for which the transcripts involved overlap mutations. Given the samples with one or more mutations *M,* and the samples with the isoform switch *S*, a Jaccard index *J* for the association of these two variables was calculated:

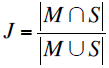

A Z-score from the Jaccard index was calculated by comparing each value *J* for an isoform switch to 100 genes with similar median isoform length. The above analysis was also repeated using only mutations that affect the protein sequence or considering the overlap with genes rather than transcript regions with similar results (see Supplementary Methods). The mutual information for the association of isoform switches and mutations, and corresponding z-score was also computed (see Supplementary Methods). The distribution of the differences between tumor and normal isoform PSIs was compared between mutated and non-mutated samples using a Mann-Whitney test. A mutual-exclusion between isoform switches and protein-affecting mutations was measured as follows: given the number of samples having an isoform switch and no mutation (*n_10_*), and those having a mutation but no isoform switch (*n_01_*), a mutual-exclusion score was defined to be:

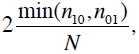

where *N* is the total number of samples. A z-score was calculated similarly as above (see Supplementary Material). For the Reactome pathway analysis the ReactomePA package from Bioconductor was used. Further details and data are provided as Supplementary Material.

## Acknowledgements

We would like to thank R. Karni, K. Hertel, Q. Morris and R. Castelo, for useful discussions. This work was supported by grants BIO2011-23920 and Consolider RNAREG (CSD2009-00080) from the Ministerio de Ciencia e Innovación of Spain and by the Sandra Ibarra Foundation for Cancer (FSI2013).

